# Reproductive success is influenced by early development: the covariation of natal body size and reproductive performance in grey seals (*Halichoerus grypus*)

**DOI:** 10.1101/2021.07.10.451928

**Authors:** Janelle J. Badger, W. Don Bowen, Cornelia E. den Heyer, Greg A. Breed

**Affiliations:** Department of Biology and Wildlife, University of Alaska Fairbanks, Fairbanks, Alaska, USA; Bedford Institute of Oceanography, Department of Fisheries and Oceans Canada, Dartmouth, Nova Scotia, Canada; Institute of Arctic Biology, University of Alaska Fairbanks, Fairbanks, Alaska, USA

## Abstract

An individual’s size when young may be an important source individual variation in lifetime reproductive performance, as size effects on ontogenetic development can have cascading physiological and behavioral consequences throughout life. Here, we explored how natal size influences subsequent reproductive performance in grey seals (*Halichoerus grypus*) using mark-recapture and repeated measures reproductive data on a marked sample of 363 females that were measured for standard length at roughly 4 weeks of age and eventually recruited to the Sable Island breeding colony. Two reproductive traits were considered: provisioning performance (mass of weaned offspring), modeled using linear mixed effects models; and reproductive frequency (the rate at which a female returns to breed), modeled using mixed-effects multistate mark-recapture models. After accounting for female age, experience, and offspring sex, we found a positive association between natal length and our measures of reproductive performance. Mothers with the longest natal lengths produced pups nearly 8 kg heavier and were 20% more likely to breed in a given year than mothers with the shortest natal lengths. Because correlation in individual body length between natal and adult life stages is weak (animals that were longer than average as pups do not always grow to be longer than average adults), the covariation between natal length and future reproductive performance reported here may act more as a carry-over effect from the size advantages afforded in the juvenile stage that allow for greater adult performance, rather than a physical trait that is maintained throughout life.

## Introduction

Life history theory predicts that maternal fitness is maximized by the reproductive strategy which results in the greatest number of offspring surviving to maturity, and subsequently producing large numbers of viable offspring themselves (Stearns 1992, Roff 1992). Variation in offspring quality may be influenced by parents through a myriad of pathways including the selection of safe and nutritious oviposition or birth sites, incubation behavior, food provisioning, defense of young, and investment in offspring size (Mousseau and Fox 1998, Krist 2011). These behaviors are costly, and reproductive strategies in long-lived species will be driven by the relationship between offspring traits and environmental conditions that determines fitness (Smith and Fretwell 1974, Mousseau and Fox 1998, Allen et al. 2008).

Offspring size is one of the most important and well-studied of these traits in evolutionary ecology, as natural selection on body size and size-related traits is ubiquitous in nature (reviewed in Sogard 1997, Krist 2011, Pettersen et al. 2015). Within species, larger offspring typically outperform their smaller conspecifics, with higher survival rates to sexual maturity (e.g. plants: Stanton 1984, marine invertebrates: Moran and Emlet 2001, Marshall et al. 2006, grey seals: Bowen et al. 2015, lizards: Sinervo 1990), enhanced resistance to starvation, environmental extremes, and predation (Sogard 1997), increased metabolic efficiency (Pettersen et al. 2015), and higher reproductive performance found in some species (arthropods: Fox and Czesak 2000, birds: reviewed in Krist 2011, marine invertebrates: Marshall and Keough 2008). Mothers may confer this advantage on their young either through a heritable genetic predisposition (possibly by choosing larger mates) or maternal effects such as nutrient transfer and protective behavior toward young (Bernardo 1996, Mousseau and Fox 1998).

The advantages of natal size are often pronounced in early stages of ontogeny, but may persist throughout life affecting reproduction and even the performance of the subsequent generation (Lindström 1999, Marshall et al. 2003, Dias and Marshall 2010). While many studies have confirmed the relationship between offspring size and survival, less is known about how the effects of natal size are subsequently manifested in adults recruited to the breeding population. Even in mammals and birds where offspring are relatively large and individuals may be tracked, estimates of the effect of an individual’s sizer when young on subsequent performance are available for only a few taxa (Clutton-Brock 1991, Festa-Bianchet et al. 2000, Fox and Czesak 2000, Crawley et al. 2017) and fewer still for natural populations. This knowledge gap is particularly apparent in long-lived iteroparous animals, where it is difficult to track individual’s reproductive performance and survival throughout adulthood. Offspring size effects on fitness would then be inaccurately estimated because key components of fitness are not measured at sufficient temporal scales (Marshall et al. 2003).

Reproductive and early life-history traits can be considered aspects of either offspring or maternal phenotype, and their evolution will therefore depend on selection operating through both offspring and maternal components of fitness (Mousseau and Fox 1998). Selection acts to maximize parental fitness, but offspring size also simultaneously influences offspring fitness. An individual’s size when young may be an important source of individual variation in lifetime reproductive performance (individual quality), as size effects on ontogenetic development can have cascading physiological and behavioral consequences throughout life (Lindström 1999). Size may mediate the expected trade-off between growth, self-maintenance, and mortality in early stages by increasing survival probabilities (avoiding starvation, predator escapement) and/or increasing foraging efficiency, allowing individuals to mature more quickly or invest in costly physiological functions to have greater lifetime reproductive output. This variation in individual quality is a key driver in natural selection and an important link between evolutionary and ecological processes (Lomnicki 1978, Cam et al. 2002, Bolnick et al. 2003, Vindenes et al. 2008, Bolnick et al. 2011, Stover et al. 2012, Gimenez et al. 2017).

The extensively studied colony of grey seals (*Halichoerus grypus*) breeding on Sable Island, Nova Scotia provides an excellent opportunity to explore the link between natal size and subsequent performance as adults. Grey seals are long-lived (*∼* 40 years), iteroparous capital breeders in which females invest heavily into the survival of a single offspring over the course of a relatively short lactation period lasting 16-18 days (Boness and James 1979, Iverson et al. 1993). During the nursing period, mothers lose a third of their body mass on average (4.1 kg per day, Mellish et al. 1999) relying only on fat reserves to produce milk and maintain metabolism, while their pups typically more than triple their birth mass (2.8 kg per day, Bowen et al. 1992). At the end of lactation, females abruptly end care and return to the sea, which conveniently allows female reproductive expenditure to be accurately measured by the energy allocated to offspring (Bowen et al. 2007). In this system, offspring size is more variable than offspring number (twins are exceedingly rare), and so size is more subject to selection for maternal fitness.

On Sable Island, grey seal pup production (a proxy for population size) has increased dramatically over the past half century with near maximum population growth of 13% per year between the 1960s and late 1990s (Bowen 2011) and a reduced rate of increase of 5-7% per year since 2004 (den Heyer et al. 2017, den Heyer et al. 2021). Associated with declines in population growth, juvenile apparent survival to reproductive recruitment has decreased by more than half from an average of 74% in cohorts born 1985-1989 to 33% in cohorts born 1998-2002 (den Heyer et al. 2013). This decline appears to be size-selective, with recent investigations finding that heavier and longer pups more likely to recruit (Bowen et al. 2015). Apparent survival to recruitment increased asymptotically with mass at weaning, but monotonically with length at weaning (Bowen et al. 2015), indicating a directional selection on skeletal size. The survival advantage of larger skeletal size may be due to increased swimming speed and agility allowing greater foraging ability and predator escapement (Sogard 1997, Hindell et al. 1999), though other physiological mechanisms cannot be ruled out. This size selection may be intensifying under density dependence, as young of the year grey seals now must make longer foraging trips and forage farther from haul-out sites than older animals which occupy foraging areas closer to rookeries (Breed et al. 2011, Breed et al. 2013), so larger-bodied animals that can swim more efficiently may experience increased survival compared to shorter conspecifics.

Here, we use a 19-year longitudinal data set of repeated reproductive measurements from individually marked, known-aged female grey seals that were measured at roughly 4 weeks of age to evaluate the influence of offspring size on reproductive success. As length is a better indicator than mass of overall skeletal size that may confer a more enduring advantage, we investigate whether variation in natal length is associated with increased reproductive performance as adults, measured using two traits: reproductive rate and offspring size at weaning. If natal length is positively associated with reproductive performance, we consider that support for a “bigger is better” hypothesis, in which maternal fitness is benefitted from bearing longer offspring that will subsequently have higher reproductive success.

## Methods

This study was conducted on Sable Island, Canada (43.93*^◦^*N, 59.91*^◦^*W), a partially vegetated sandbar on the Scotian Shelf roughly 160 km off the coast of Nova Scotia, Canada, during the 1989-2020 breeding seasons. The breeding season at this colony spans early December through early February, with 91.2% of pups born by mid-January (Bowen et al. 2007, den Heyer et al. 2021). Sable Island supports the largest breeding colony of grey seals in the world with an estimated 87,500 pups (SE = 15,100) born on the island in 2016, comprising 80% of the total grey seal pup production in the Northwest Atlantic (den Heyer et al. 2021).

### Data Collection

Our 19-year study (2002-2020) was conducted on a subset of female grey seals born on Sable Island from 1998-2002 that survived to recruit to the breeding colony, as part of a larger program led by the Department of Fisheries and Oceans, Canada (DFO). Individuals were marked at roughly 4 weeks old, shortly after weaning, with unique alpha-numeric hot-iron brands in each year 1998-2002. Prior to marking, researchers recorded standard dorsal body length (to the nearest cm) of these individuals while they were sedated with diazepam (*∼* 0.4 mg/kg body mass, Sandoz Canada, Boucherville, Quebec, Canada) to allow for accurate measurement (Bowen et al. 2015). These permanent brands allowed reliable identification of individuals over the course of their lives. Females can recruit to the breeding population as early as 4 years old, but this is uncommon, and the average age of first reproduction is 6.5 ± 0.21 SE years for these cohorts (den Heyer et al. 2013) with 87% of females recruited at or before age 7 (Bowen et al. 2015). Each breeding season since 2002, teams of researchers conducted 5-7 roughly weekly censuses of branded females returning to the island to give birth and mate. Once sighted, branded individuals with dependent pups were visited daily but generally not disturbed. Prior to weaning, pups were sexed and marked with semipermanent, uniquely numbered tags in the hind flipper to ensure accurate identification after the marked female ended lactation and returned to sea, leaving her pup in the colony. Females attend their pups continuously throughout lactation. Therefore, once a pup was sighted alone, it was considered weaned and weighed to the nearest 0.5-kg.

The probability of observing a marked female during any given year includes both the probability the female is present, and the probability that she is detected given presence at the breeding colony. A recent analysis of this population indicated that, if a female rears a pup on the island, there is less than a 5% chance researchers will fail to detect her in at least one resighting census (Badger et al. 2020). Individuals that are not rearing pups can be skittish and may flee to the water, resulting in a lower sighting probability than females nursing and defending young. Grey seals are highly site philopatric, and once recruited to a breeding colony, will very rarely pup elsewhere (Bowen et al. 2015). Thus, we are able to reliably follow the reproductive history of individuals, and do not expect permanent emigration to other colonies to be significant source of sighting error.

Individual sighting histories were collected from age at first reproduction (first sighting in breeding colony) until the most recent year of our study, 2020. Sighting histories of individuals were scored as a 0 (not sighted) or 1 (sighted) for each year 2002 to 2020. Females sighted in only one breeding season were omitted from this analysis to ensure that they had in fact recruited to the Sable Island breeding population and we have adequate data to estimate reproductive performance.

### Statistical Analysis

In this analysis, we were interested in understanding how a female’s size during early life stages influences subsequent reproductive success once she has matured. To do this, we analyzed the effect of natal length (*L_M_*, after weaning, but prior to independent foraging) on her reproductive performance, measured two ways: annual provisioning performance and reproductive frequency (both described below). We used generalized mixed-effect additive and linear models to determine the effect of maternal natal length *L_M_* on these traits, and accounted for imperfect detection in reproductive rate using a multi-state capture-recapture model in a Bayesian framework (Gimenez et al. 2007, Lebreton et al. 2009, Kéry and Schaub 2012).

#### Modeling annual provisioning performance

During lactation, grey seal pups consume only milk provided by the female, and as capital breeders, females fast for the entire lactation period and provision pups exclusively from energy stores. Therefore, in our study, the body mass of a pup at weaning is a reasonable estimate of the energy (i.e. nutrients) transferred to young, and is of critical importance for pup survival (Hall et al. 2001, Bowen et al. 2015). We modeled the weaning mass of pup *j* born to female *i* in year *t* (*mass_j,t_*) as a linear mixed-effects model with female experience (parity, i.e. *par*; because this effect tends to plateau, it was discretized into 1, 2, and 3+ parities), offspring sex, and a quadratic effect of standardized female age as covariates along with random individual and year intercepts:

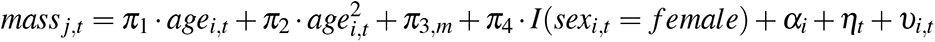

Where linear parameters are held in the vector ***π*** = {*π*_1_, *π*_2_, *π*_3_, *π*_4_} and represent linear and quadratic age effects, effect of female experience, and pup sex, respectively; and **I** signifies an indicator variable, and *m* denotes the parity group (1, 2, or 3+) of female *i* in year *t* so *m* ∈ {1, 2, 3}. *α_i_* is the random effect of individual such that 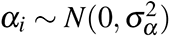, *η_t_* reflects the random year effect, where 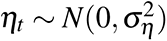, and *υ_i,t_* is the error term where 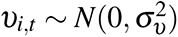.

We tested the effect of natal length *L_M_* on the history of her pup weaning masses comparing this null model to models including *L_M_* as a linear term and a quadratic term (Table 1). We also included a model in which the effect of *L_M_* on offspring size varies with parity, such that the effect may diminish over time (Dias and Marshall 2010). Models were fit using the `lmer` function in package `lme4` (Bates et al. 2015), and support for model configurations was determined via likelihood ratio tests using the `anova` function offered in `R` (R Core Team 2020).

**Table 1:** Four competing linear mixed effects models to describe the effect of natal length on her reproductive performance, measured as offspring mass.

| Model | Form | AIC | LRT $p$ value |
| --- | --- | --- | --- |
| Mod 0: Null | $mass_{j,t} = \pi_1 \cdot age_{i,t} + \pi_2 \cdot age_{i,t}^2 + \pi_{3,m} + \pi_4 \cdot I(sex_{i,t} = female) + \alpha_i + \eta_t + v_{i,t}$ | 17713 | - |
| Mod 1: Linear effect of natal length | Mod 0 + $\pi_5 \cdot L_{M,i}$ | 17702 | $p < 0.001$ |
| Mod 2: Quadratic effect of natal length | Mod 0 + $\pi_5 \cdot L_{M,i} + \pi_6 \cdot L_{M,i}^2$ | 17702 | $p = 0.176$ |
| Mod 3: Interactive effect with maternal experience | Mod 0 - $\pi_{3,m} + \pi_5 \cdot L_{M,i} + \pi_6 \cdot L_{M,i} \cdot I(par_{i,t} = 2) + \pi_7 \cdot L_{M,i} \cdot I(par_{i,t} = 3)$ | 17705 | $p = 0.602$ |
| Mod 4: Cohort effects | Mod 0 + $\pi_5 \cdot L_{M,i} + \pi_c$, where $c \in \{1998, 1999, 2000, 2001, 2002\}$ | 17707 | $p = 0.631$ |

Where $mass_{i,t}$ is the mass of the weaned pup born to female $i$ in year $t$ . Parameters $\pi = \{\pi_1, \pi_2, \pi_3, \pi_4\}$ reflect the quadratic age effect, effect of female experience, and pup sex, respectively, and $\pi \in \{\pi_5, \pi_6, \pi_7\}$ describe the effect of maternal natal length, $L_M$ under different models. $\alpha_i$ is the random effect of individual such that $\alpha_i \sim N(0, \sigma_\alpha^2)$ , $\eta_t$ reflects the random year effect, where $\eta_t \sim N(0, \sigma_\eta^2)$ .

#### Modeling reproductive rate

The second reproductive trait, reproductive rate, is defined as the probability an individual will return to the island to give birth in any given year. We estimated the effect of a female’s natal length *L_M_* on her reproductive rate by modeling her reproductive history as a Markov chain in a multi-state capture re-capture modeling framework. Between her first and last sightings on the island during our study, a female transitions among three reproductive states: initially a first time breeder *F*, then switching between a breeder state *B*, or non-breeder state *N*. Reproductive frequency is then defined as the probability of transition *ψ^kB^* into a breeding state *B* from any reproductive state *k*. An individual’s state transition from year *t* to *t* + 1 is modeled as a categorical trial with probabilities of transition *ψ^ks^* from state *k* to state *s*. We used mixed-effects logistic regression embedded in this multistate model to account for standardized female age, previous breeding state, and random individual and year effects in probability of breeding (*ψ^kB^*):

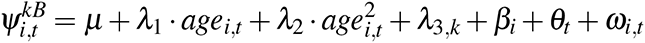

Where parameters ***λ*** = {*λ*_1_, *λ*_2_, *λ*_3_} represent the quadratic age effect and the effects of the previous breeding state *k*, respectively, where **I** denotes an indicator variable and parameters *λ*_3_*_,k_* sum to zero. *β_i_* is the random effect of individual such that 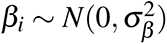, *θ_t_* reflects the random year effect, where 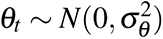, and *ω_i,t_* is the error term where 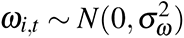.

Similar to above, we tested the effect of natal length *L_M_* on her reproductive rate by comparing this null model to models including *L_M_* as a linear term and a quadratic term (Table 2). Further, we included a model in which the effect of *L_M_* on offspring size varies with parity, such that the effect may diminish over time.

**Table 2:**
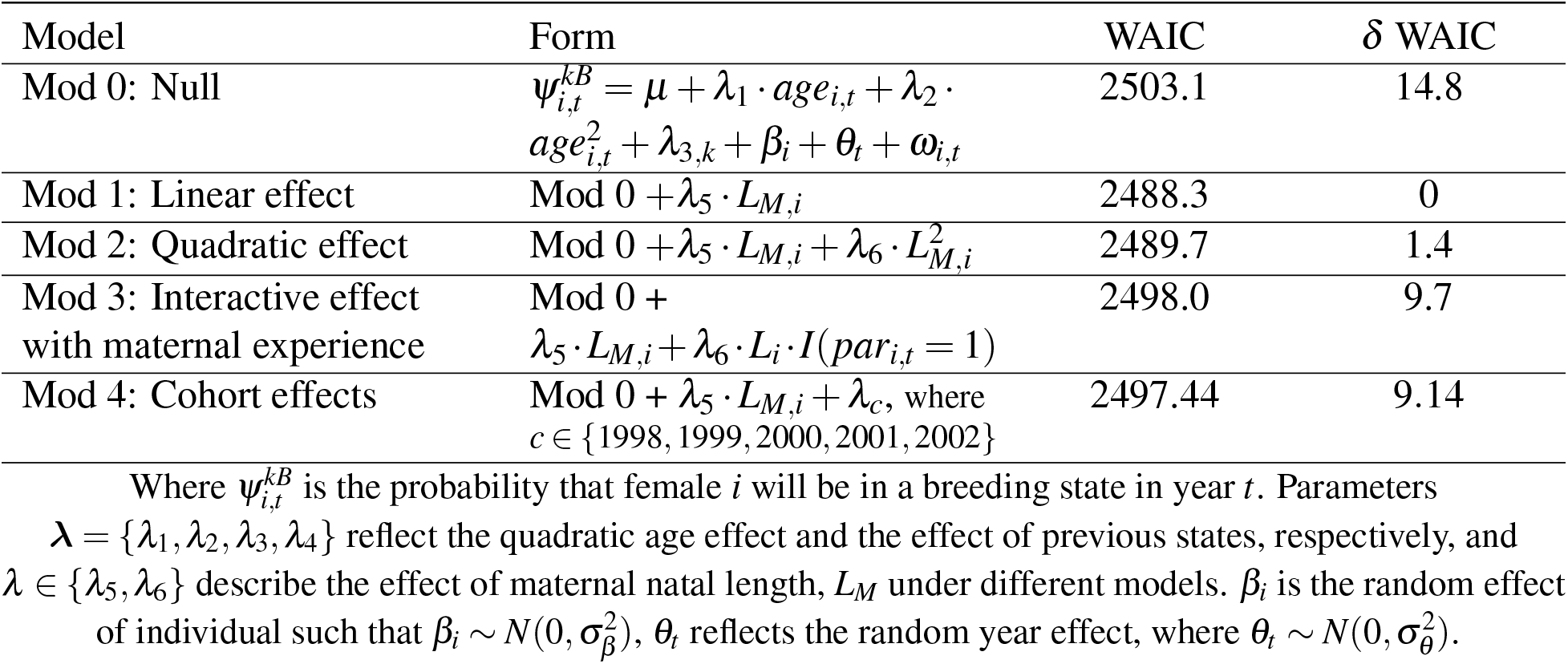
Four competing multistate mixed effects mark-recapture models to describe the effect of natal length on her reproductive performance, measured as reproductive rate.

Multistate models can also be used to detect a cost of reproduction (e.g. Beauplet et al. 2006, Hernández-Matías et al. 2011, Chambert et al. 2013, Stoelting et al. 2015, Johns et al. 2018, Badger et al. 2020). A common approach is to determine whether breeding at time *t* negatively affects an individual’s probability of surviving from time *t* to *t* + 1 or its probability of breeding at time *t* + 1. In the model used here, one way in which a cost of reproduction may be observed as a higher probability of transition *ψ* into a breeding state *B* from a nonreproductive state *N*, i.e. *ψ^NB^* > *ψ^BB^*.

A Bayesian approach was used for estimation and implemented in the software program JAGS 4.2.0 using the `R` interface `rjags` (Plummer 2003, R Core Team 2020, Plummer 2018). Parameters ***λ*** were assigned diffuse normal prior distributions *N*(0, 1000). Random year term *θ* was specified hierarchically following a normal distribution, 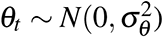, and individual terms *β_i_* were pulled from a 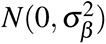. We specified a Unif(0,10) prior for *σ_θ_* and *σ_β_*.

Markov chain Monte Carlo (MCMC) methods were used to sample the posterior distributions of the parameters of interest. For each of the competing models, we ran three chains in parallel using package `dclone` (Solymos 2010) with different sets of initial values. The first 10,000 MCMC samples were discarded, known as the burn-in period, after having checked that convergence was satisfactory. Convergence was visually assessed using sample path plots in conjunction with the Brooks-Gelman-Rubin diagnostic 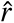 (Brooks and Gelman 1998), with values close to 1.00 indicating adequate convergence. Chains then ran for 100,000 iterations after burn-in, and a total of 3,000 MCMC samples (every 100th sample of each chain) were used for inference. We determined that a covariate had an effect if a 95% credible interval (CRI) of the posterior distribution of that parameter did not include 0. We assessed support for inclusion of natal length using a measure of out-of-sample predictive ability of each model, the Widely Applicable Information Criterion (WAIC, Watanabe 2010), where a model with a smaller WAIC is judged a better fit.

## Results

We analyzed the reproductive histories of 363 females born from 1998-2002 that gave birth to a total of 3457 pups. 2.5% (9/363) of those females recruited to the breeding population at age 4, 31.4% (114/363) had their first birth at the age of 5, 24.5% (89/363) at the age of 6, and 30.5% (111/363) recruited after age 6. From primiparity to the most recent year of the study, 2020, females had an average of 10 pups (*SE* = 4.48, ranging 1 to 17). These female’s lengths during early development (*L_M_*, i.e. at around 4 weeks of age) ranged from 90-123 cm, with an average of 110.7 cm (*SE*= 4.28). We did find a cohort effect on *L_M_* (ANOVA, p = 0.003), where females born in 2002 that recruited to the breeding population were significantly longer (Tukey HSD, Figure 4).

### Effect of natal length on future reproductive performance

A female’s natal length *L_M_* was positively associated with her future provisioning performance (*p* < 0.001, Table 1). The best supported model describing pup weaning masses included an additive, linear effect of natal length as a covariate, though there was some support for a quadratic effect (Table 1, Appendix B: Table B2). The longest females (when young) proceeded to give birth to offspring that weaned 7.97 kg heavier, on average, than their shortest conspecifics (Table 3). Though we expected natal body length to have a greater effect on early parities (such that the effect weakened over time), we found no support for an interactive model between *L_M_* and parity (Table 1, Appendix B: Table B3). Repeatable differences among individuals accounted for 41% of the variance in pup weaning mass. Year accounted for only 10.8% of the variance in weaning mass, suggesting that among year environmental effects were small. Natal length was also positively associated with a female’s future reproductive rate. Model output from fitted multistate Markov models estimated that natal length accounts for the spread in annual reproductive probability to range from 0.715 for the shortest females to 0.916 for the longest (Table 4, Figure 2). Model fits displayed no evidence of inadequate convergence to stationary distributions.

**Figure 1:**
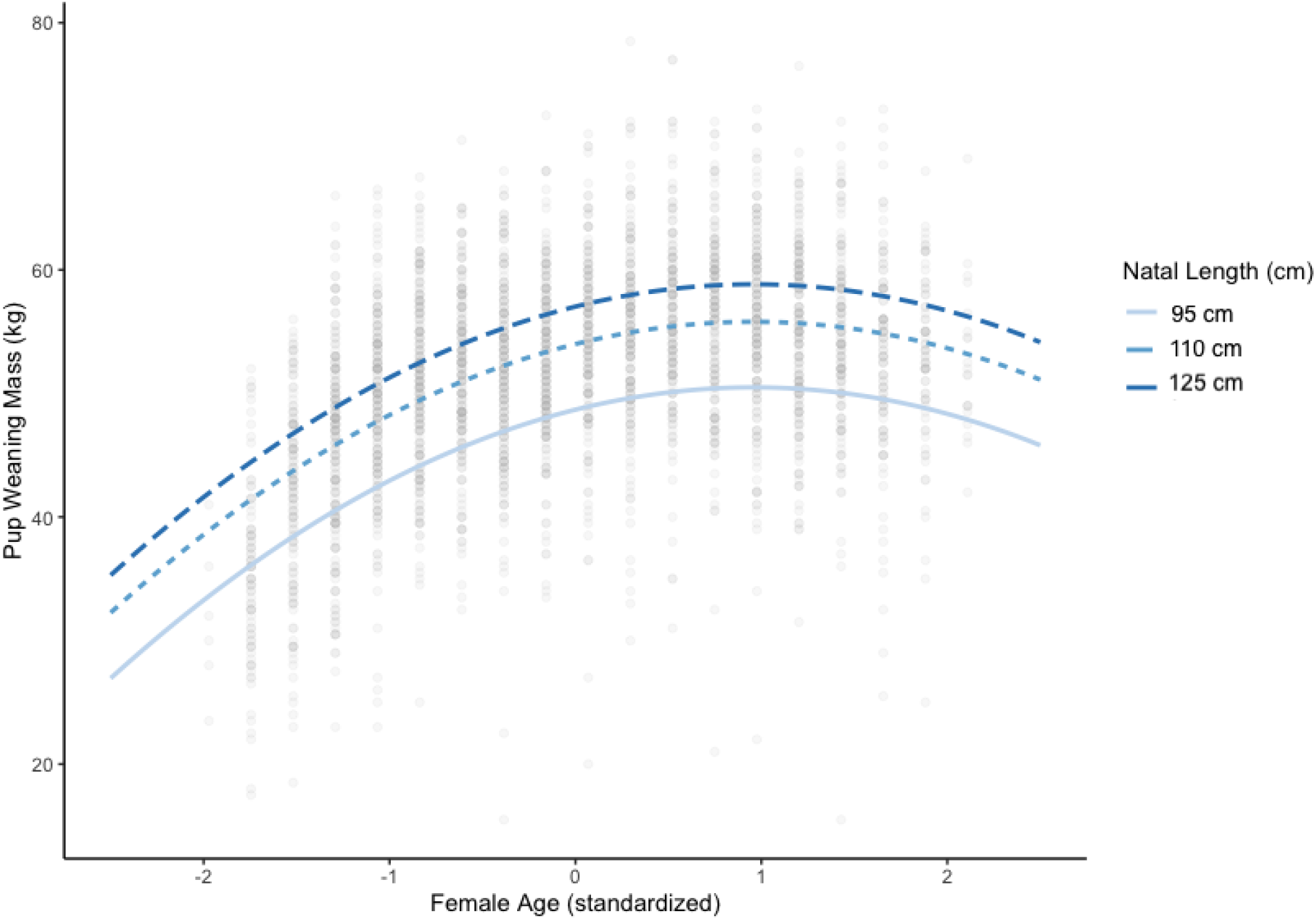
The estimated effect of natal length on provisioning performance as a female ages. Lines are 0.025%, 50%, and 97.5% quantiles of natal lengths corresponding to 95 cm, 110 cm, and 125 cm.

**Figure 2:**
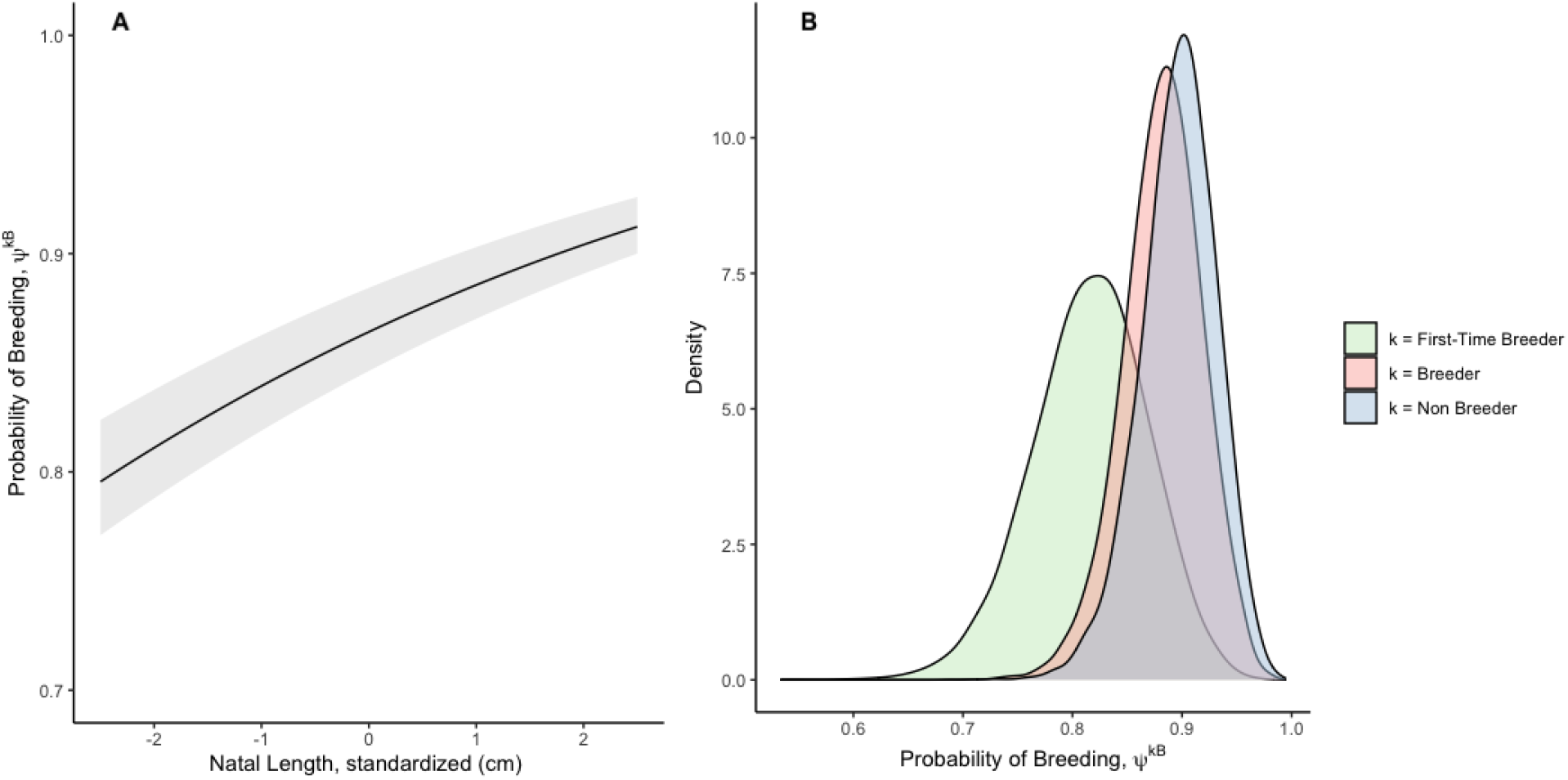
Results from the Markov chain multi-state model describing probability of breeding, *ψ^kB^*, as a function of (A) natal length, and (B) the female’s previous state in year *t*-1.

**Table 3:** Parameter estimates for favored linear mixed effects model describing variation in pup weaning mass as a function of maternal age, experience (parity), pup sex, natal length *L_M_*, and random effects of year and individual.

| Parameter | Mean | St. Error |
| --- | --- | --- |
| Intercept | 49.22 | 0.791 |
| $\pi_1$ | 14.46 | 1.26 |
| $\pi_2$ | -11.88 | 1.19 |
| $\pi_3 \vert par_{i,t} = 2$ | 4.01 | 0.55 |
| $\pi_3 \vert par_{i,t} = 3$ | 6.23 | 0.63 |
| $\pi_4$ | -2.26 | 0.22 |
| $\pi_5$ | 1.07 | 0.29 |
| $\sigma_{ID}^2$ | 4.67 | |
| $\sigma_{year}^2$ | 1.29 | |
| $\sigma_{residual}^2$ | 5.45 | |

**Table 4:** Posterior mean, SD, 2.5%, 50%, and 97.5% quantiles, and convergence diagnostic 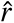 of parameters for preferred multistate model, describing variation in reproductive rate 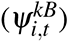 as a function of previous reproductive state, quadratic effect of maternal age (*λ*_1_, *λ*_2_), linear maternal length as young *L_M_* (*λ*_5_), and random effects of individual and year. The effect of previous state is reported here as transition rates among F, B, and N for ease of interpretation.

| Parameter | $\hat{r}$ | Mean | SD | 2.5% | 50% | 97.5% |
| --- | --- | --- | --- | --- | --- | --- |
| $\psi_{i,t}^{BB}$ | 1.003 | 0.861 | 0.038 | 0.784 | 0.861 | 0.935 |
| $\psi_{i,t}^{FB}$ | 1.003 | 0.779 | 0.056 | 0.665 | 0.780 | 0.888 |
| $\psi_{i,t}^{NB}$ | 1.003 | 0.878 | 0.037 | 0.803 | 0.879 | 0.950 |
| $\lambda_1$ | 1.016 | 0.385 | 0.083 | 0.125 | 0.412 | 0.463 |
| $\lambda_2$ | 1.016 | -0.355 | 0.083 | -0.440 | -0.380 | -0.096 |
| $\lambda_5$ | 1.001 | 0.549 | 0.020 | 0.513 | 0.549 | 0.594 |
| $p$ | 1.007 | 0.975 | 0.021 | 0.924 | 0.980 | 0.999 |
| $\sigma_\beta^2$ | 1.006 | 0.895 | 0.151 | 0.673 | 0.869 | 1.269 |
| $\sigma_\theta^2$ | 1.001 | 1.310 | 0.695 | 0.371 | 1.169 | 3.036 |

### Cost of reproduction in breeding rate

In this analysis fit to the reproductive data of individuals from the 1998-2002 cohorts, the fitted multistate model estimated somewhat (*∼* 2%) higher reproductive probabilities for individuals that did not breed in the previous year (Table **??**). However, previous analyses on a larger subset of this population including individuals born in the 1960s - 1980s, did not find evidence for a cost of reproduction expressed in reproductive rate. In one of these previous analyses, individuals that reproduced in the current year were on average 11% more likely to breed the next year than those that skipped reproduction (Badger et al. 2020, den Heyer and Bowen 2017, Figure 3). Importantly, females born in the 1960s-1980s recruited during a period of exponential growth with population densities much lower than the females recruiting in the present study (den Heyer and Bowen 2017). The result of this current analysis, indicating a slight cost under higher population densities, contrasting with the previous studies indicating no cost when population densities were lower suggest that the cost of reproduction may only be expressed at higher population densities.

**Figure 3:**
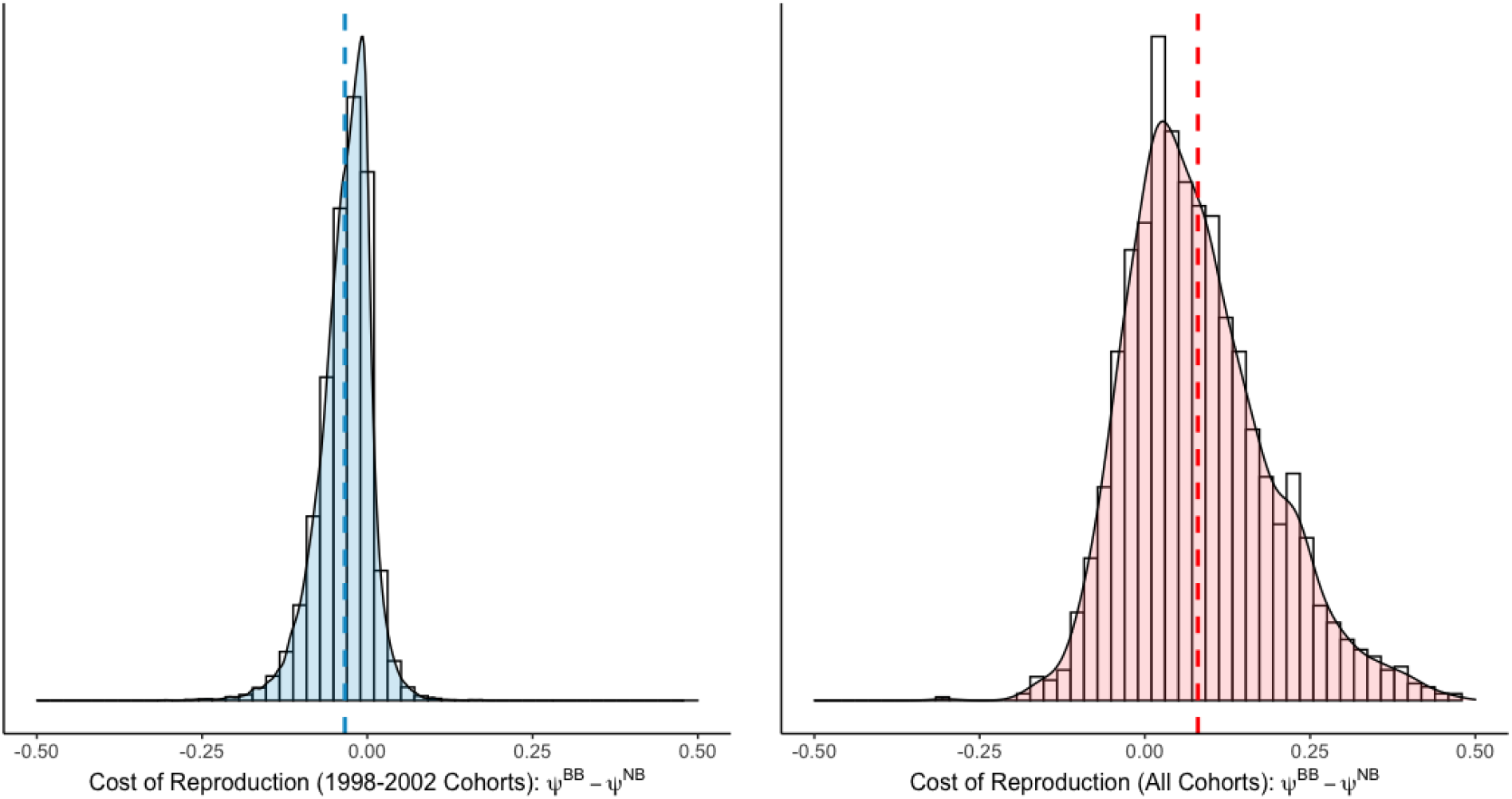
The cost of reproduction is estimated by finding the difference between reproductive probabilities of non-breeders and breeders: panels depict posterior distribution of *ψ^BB^* minus posterior distribution of *ψ^NB^* for (A) output of the models reported here, estimating reproductive probabilities for females born from 1998-2002, and (B) the output from Badger et al. 2020, a similar model estimating reproductive probabilities for females born 1962, 1969, 1970, 1973, 1974, 1985-87, 1989, and 1998-2002. Note that for (B), the models did not estimate a cost of reproduction in terms of reproductive rate, where *ψ^BB^* > *ψ^NB^*, i.e. current reproduction does not incur a “penalty” to future reproduction. By contrast, our sample of females (A) show a slight cost of reproduction *ψ^BB^* < *ψ^NB^*, where individuals are slightly more likely to breed in a given year if they had skipped reproduction previously.

**Figure 4:**
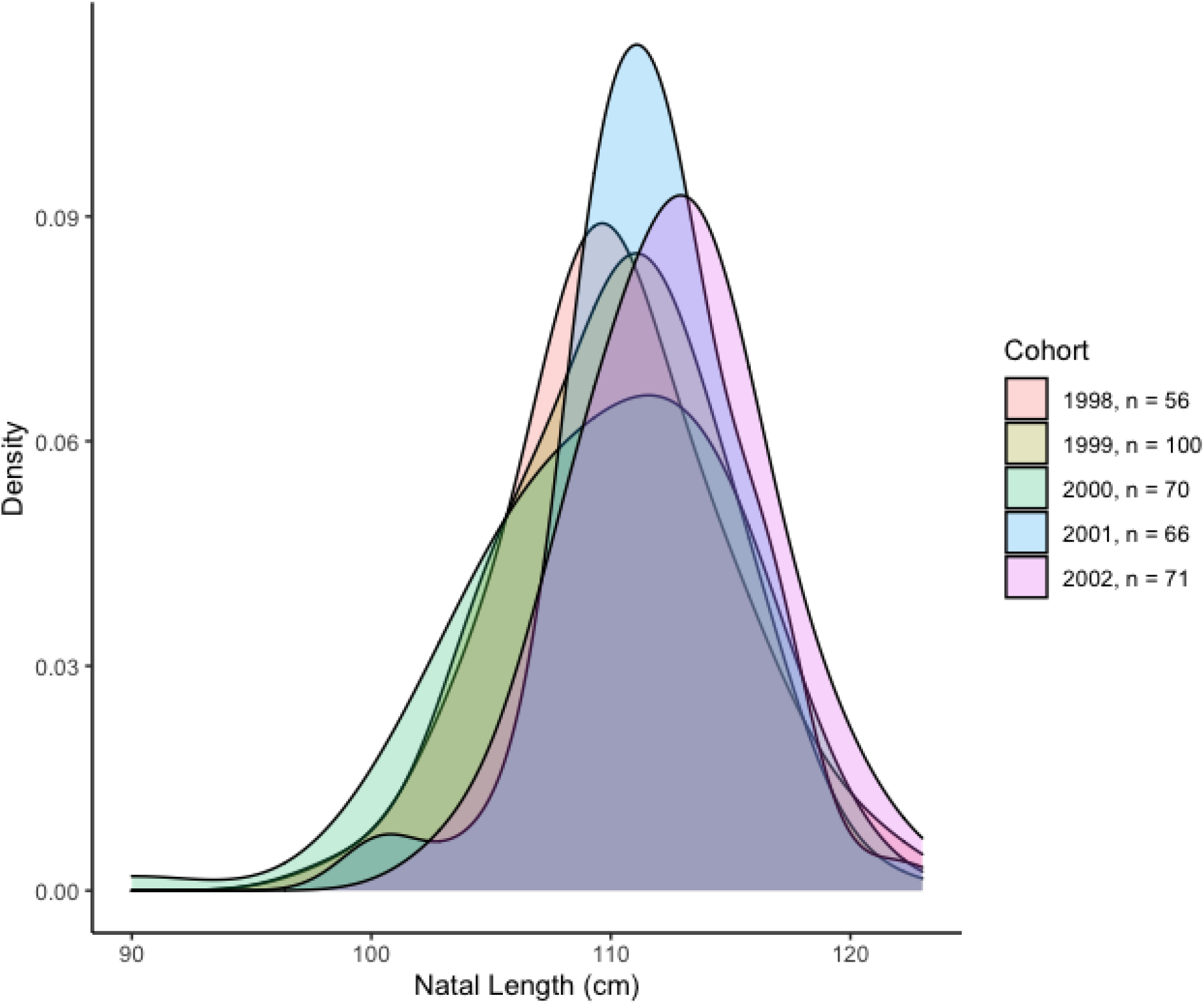
Density plots of the distribution of natal lengths of our sample of females by cohort, 1998-2002.

**Figure 5:**
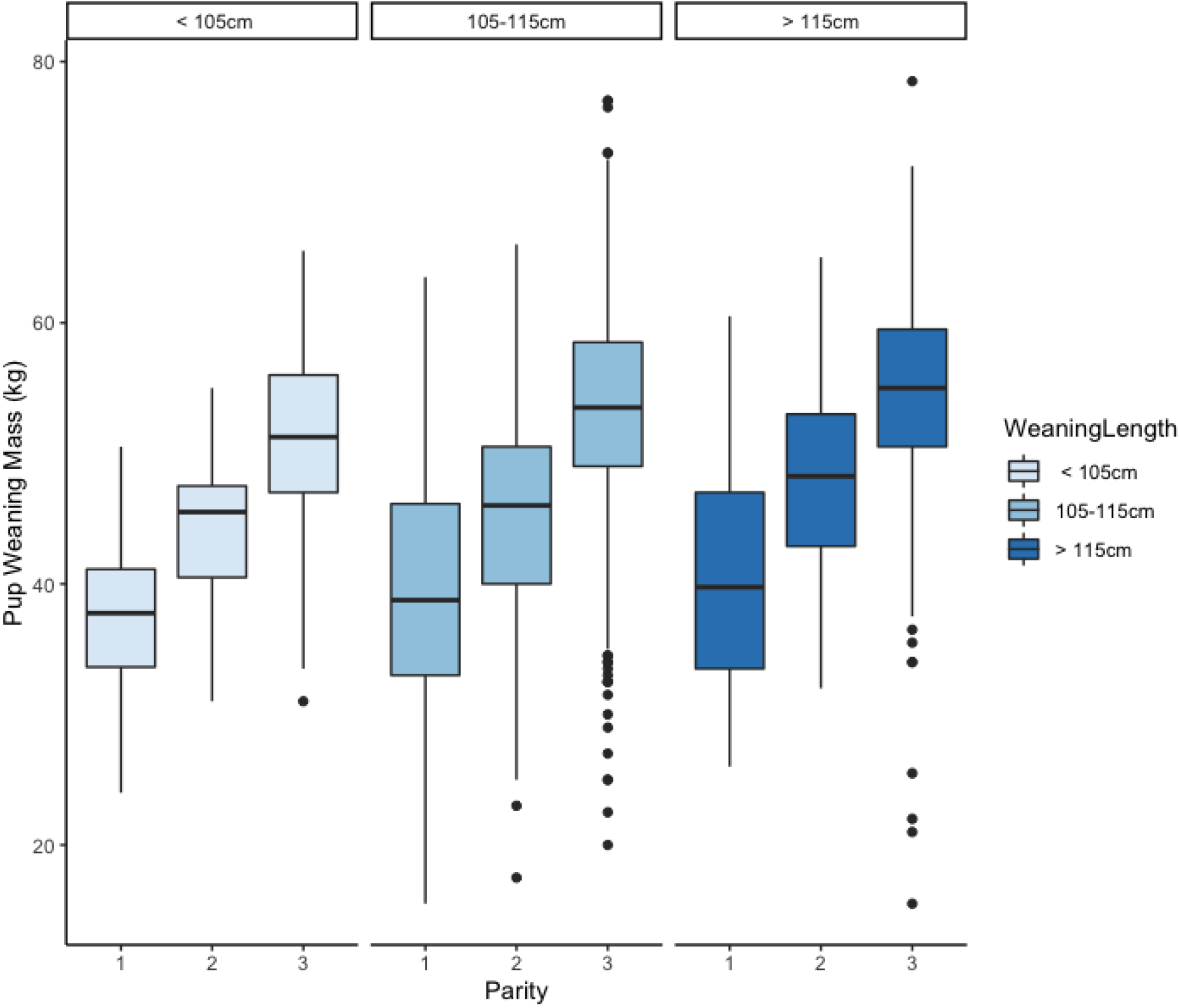
There is no evidence for an interactive effect of natal length and parity–effect of natal length on pup weaning mass does not taper off (*p* > 0.05, Table 1). Boxplots of pup weaning masses for individuals with short (90 - 105 cm), average (105 cm - 115 cm), and tall (115 - 125 cm) natal lengths (panels) over the 1st, 2nd, and 3+ parities.

### Cohort effects in reproductive performance

After detecting a cohort effect in maternal natal length (*L_M_*), we incorporated cohort effects into reproductive performance models of breeding rate and offspring mass. Individuals from the 2002 cohorts had lighter pups on average than individuals from other cohorts (Appendix B: Table B4), though this model performed worse in out-of-sample predictive accuracy than models without cohort effects (Table 1). In reproductive rate, our multistate models also estimated a lower breeding rate of individuals born in the 2002 cohort (Appendix B: Table B4). However, this model also performed poorly relative to models not including cohort as a covariate (Table 2). We further caution the interpretation of the result of this multistate model including cohort effects as we were not able to control for the effects of maternal age due to issues with convergence likely stemming from multicollinearity of the age, cohort, and parity variables.

## Discussion

Here, we found positive covariation between an individual’s length during early ontogeny and their reproductive performance, measured by two traits, using nearly two decades of longitudinal reproductive data on a large sample of female grey seals. Females that were longer when young were more likely to reproduce in any given year and had larger pups than their shorter conspecifics. This result is consistent with a “bigger is better” hypothesis (Bowen et al. 2006), in which longer offspring mature to have higher reproductive success as adults.

Mothers with the longest natal lengths produced pups nearly 8 kg heavier and were 20% more likely to breed in a given year than mothers with the shortest natal lengths. This spread in offspring size and reproductive frequency can drive substantial variation in lifetime reproductive output. Badger et al. (2020) found that reproductive frequency and the probability of weaning a pup large enough to survive independently were well correlated among individual grey seal females, and over their lifetimes higher performing females will produce 1.83 times more successful pups than poorer performers. Further, below population average weaning mass (51.5 kg), pup survival is dependent on mass (Hall et al. 2001, Bowen et al. 2015), and an 8 kg spread can have a large impact on the probability a female’s offspring will mature to reproduce. Bowen et al. (2015) estimated that below 51.5 kg, each 1 kg decrease in pup weaning mass corresponded to a 0.12 decrease in survival to reproductive recruitment (on the logit scale). Consequently, the results we report here have important implications for both maternal and offspring fitness.

Body mass may be more convenient to measure in many systems because body posture can greatly impact standard length measurements. For Sable Island grey seals, standard length is only taken when pups are sedated in preparation to be marked, which only occurs in certain sampling years (1998-2002 and 2014-2016). So, in this analysis weaning mass, not weaning length, was used as a proxy of reproductive success even though positive effects of offspring weaning mass on offspring fitness attenuate at heavier weights (Bowen et al. 2015). This analysis could be strengthened with further data on offspring length coupled with mass to understand the parameters of reproductive success.

### Implications for maternal fitness

A female’s fitness does not simply depend on the number of offspring she produces, but also on the reproductive success of those offspring, which will ultimately determine her genetic contribution to the population through subsequent generations. We show that grey seals mothers increase maternal fitness by producing longer pups, as longer pups mature to be more productive mothers. Thus larger skeletal size of offspring may boost maternal fitness by favoring the propagation potential of her genetic material.

While the effect of offspring body size on maternal fitness has been extensively studied (Lack 1947, Cody 1966, Smith and Fretwell 1974, Stearns 2000, Krist 2011, Rollinson and Hutchings 2013, Pettersen et al. 2015), the focus on body length or skeletal size rather than total mass or fat content/energy stores is less common. Body length and fat content of offspring reflect different aspects of maternal quality: innate genetic material versus maternal effort and investment. Producing fat offspring may result in significant reproductive energy expenditure, in both acquisition of resources (e.g. foraging efficiency, prey choice, instraspecific competition) and effectively transferring energy resources to offspring (e.g. lactation efficiency, nursing behavior). Increasing an offspring’s body length, however, has a much larger genetic basis–one or both of the parents must carry genetic material that may be expressed to promote skeletal size in their offspring. The genetic architecture of body size is unknown for pinnipeds, but divergent selection experiments in domestic mammal species suggest that dozens to thousands of loci underlie variation in body size (Kemper et al. 2012).

Allocation theory predicts an asymptotic relationship between offspring size and offspring survival because parents are expected to receive decreasing returns on investment in offspring fitness after a certain point (Smith and Fretwell 1974). Previous analyses into this population suggest a stabilizing selection on body mass at weaning near the observed average, where offspring survival to recruitment levels and eventually decreases with increasing mass (Bowen et al. 2015). However, body length appears to be subject to directional selection (at least in this ecological environment), evidenced by a monotonically increasing relationship between body length and offspring survival to reproductive recruitment (Bowen et al. 2015). Early growth rate will vary among individuals owing to their genetic makeup and the individual’s foraging success under differing environmental conditions (Madsen and Shine 2000, Harrison et al. 2011). While fat mass provides crucial resources during the transition to independent foraging, fatter pups are likely more buoyant, and therefore less efficient foragers and more vulnerable to predation (Sogard 1997, Hindell et al. 1999). Young of the year grey seals do make longer foraging trips and forage farther from haul-out sites than older animals (Breed et al. 2013), which suggests lower foraging efficiency relative to adults. Longer individuals, however, may gain a tangible benefit throughout early stages due to greater swimming speeds and outgrowing predators. The possible mechanisms driving relationships between early traits and survival remain to be tested, but results from this analysis indicate benefits of length have a persistent effect on fitness and potential for strong transgenerational effects on reproductive output in this species.

### Natal length as a source of individual variation in quality

Recent analyses into this population support the presence of substantial differences in quality (i.e. lifetime reproductive success) among individuals. Though it is expected that maternal effects on offspring size are most significant at early life history stages (Dias and Marshall 2010), with compensatory growth or other factors reducing impact later in life (e.g., domestic sheep, Wilson and Réale 2006, red squirrels, Wauters et al. 1993), our results suggest variation in natal body length may partially explain some of the observed variation in individual quality across an individual’s lifetime. Individuals that were longer during early ontogeny consistently outperform those of shorter lengths in both survival to recruitment to the breeding population (Bowen et al. 2015) and reproductive success once recruited (this study).

As capital breeders that rely heavily on fat reserves to support lactation, maternal body fat stores are likely more influential to reproductive performance than adult body length. However, adequate fat reserves depend on effective acquisition and conservation of food energy, so traits affecting foraging such as body length could drive substantial variation in reproductive success. Foraging is a complex suite of behaviors that entails energetic costs associated with supporting basal metabolism, locomotion, and the digestion of prey. Although larger animals have higher absolute metabolic requirements than smaller ones, larger individuals exhibit lower mass-specific rates of metabolism which confers a suite of physiological and ecological benefits at greater body sizes (Kleiber 1947, Glazier 2005, Gearty et al. 2018). These advantages include a low cost of transport, enhanced fasting ability, and, for animals such as seals, the ability to dive longer and deeper affects the rate and ability to capture prey (Peters 1983, Costa 1993).

However, the extent to which body length, independent of body mass, may offset the energetic cost of foraging is unknown in many systems, including grey seals. In Weddell seals, Wheatley et al. (2006) found that postpartum mass of shorter females was significantly lower in years of poor environmental conditions whereas the mass of longer females did not differ between years. This suggested that shorter females were less successful foragers than their larger conspecifics and may generally be more susceptible to environmental variation (Wheatley et al. 2006). If longer females are more successful foragers, or more robust to environmental variation, they would have a distinct advantage in accumulating and storing energy needed for reproduction. Inclusion of female body mass as a covariate in these analyses could help determine whether maternal energy stores, independent of body length, help predict the variation in individual life histories, but these data were not available on a large scale.

Alternatively, length may be advantageous in growing juvenile stages for grey seals, but the effect of this trait could attenuate over time. Large skeletal size as an adult could alternatively be subject to stabilizing selection as larger individuals experience different physical constraints and energetic costs in their environment (Williams et al. 2000). Increasing body size will increase costs to sustaining body condition and maintaining buoyancy in the water column. Though smaller animals have a higher mass-specific metabolism, their absolute food energy requirements are lower (Peters 1983, Costa 1993) and so could be less vulnerable to food scarcities. Smaller prey items are relatively unprofitable to larger individuals than smaller individuals, requiring additional costly prey captures to reach energy requirements, decreasing the efficiency of a foraging bout (Costa 1993) and competitive ability under resource limitation (Clutton-Brock 1988). This may be of special importance in the Northwest Atlantic, where size-selective fishing and regime shifts (Frank et al. 2005) have resulted in many fish species maturing earlier and smaller (Swain et al. 2007), releasing of smaller pelagic fishes from predation, and major alteration of the overall size distribution of available prey. Length may have further implications for physiological processes such as circulation and thermoregulation–longer body lengths at similar masses will have larger surface area to volume ratios, increasing heat loss and thus energetic expenditure. Grey seals that breed on North Rona, Scotland behaviorally thermoregulate to avoid thermal stress during the breeding season (Twiss et al. 2002), though the North Rona colony uses warmer autumnal months as their breeding season.

While our study only followed female offspring through early adulthood, male pups likely also benefit from increased skeletal size later in life. In big horn sheep, *Ovis canadensis*, another mammalian species with male-male intrasexual selection, early size explained more of the variation in adult performance in males than for females, and females were better able to compensate for small natal size than males (Festa-Bianchet et al. 2000). In grey seals, body length was found to be an important factor in tenure (time spent at a site among females) of adult males, increasing access to females (Lidgard et al. 2012). Longer males may have greater competitive ability and endurance (Murphy 1998), and thereby be more successful at defending harems through intimidation displays and male combat. However, no strong relationship with reproductive success was found, as measured by observed number of mates, which may be due to difficulty in measuring reproductive success of male grey seals (Lidgard et al. 2012). Male body mass explained more variation than body length in reproductive success, but as a quadratic relationship with the number of consorts mated and the estimated number of pups sired such that males of intermediate masses experienced the greatest success (Lidgard et al. 2005).

Regardless, longer pups do not necessarily mature into longer adults. Bowen et al. (2015) found a positive, but weak correlation between body length of these female pups and their length at primiparity (age at first reproduction), and length data collected sporadically since suggests this relationship is small through adulthood. Natal body length accounted for 6% of the variation in primiparous length (n = 325, Bowen et al. 2015), 4.6% of variation in body length of adult females during early adulthood (5-10 yrs, n = 268, unpublished data) and 4.3% of the variation in body length of older females (10+ years, n = 29, unpublished data). So, it is unlikely that the results reported here are solely due to longer individuals when young remaining long throughout life. Growth and reproduction are involved in a classic physiological trade-off, and further somatic investment during reproductive years may be unfavorable to maximizing fitness (Partridge and Harvey 1988, Green and Rothstein 1991,Stearns 1992), though this relationship may be mediated by individual quality (Clutton-Brock 1984,van Noordwijk and de Jong 1986).

### Evolutionary considerations of natal body length and reproductive performance

As long-lived iteroparous mammals that must allocate their reproductive effort over many years to maximize fitness, the possession of a “longer natal” genotype may provide an edge to produce pups that are more likely to survive to reproduction, and have greater reproductive success, producing larger pups themselves. Across a wide variety of taxa, larger mothers produce larger offspring (reviewed in Lim et al. 2014, Cameron et al. 2016). Length, if a heritable trait that confers a reproductive advantage, could further ensure future generations to also have higher reproductive probabilities. Heritable individual quality has recently attracted growing interest among ecologists (Chambert et al. 2014, Cam et al. 2016, Gimenez et al. 2017), but few studies have identified a source or specific traits contributing to heritable phenotypic quality (Gimenez et al. 2017). Significant heritability of length traits have been estimated in many systems (e.g. hindleg length Soay sheep, Wilson et al. 2007) though the extent of heritability in body length in grey seals (and indeed seals and marine mammals generally) has not been tested.

In the grey seal cohorts we studied, there is evidence for positive selection for natal length in recruitment (Bowen et al. 2015), and, the results reported here indicate that natal length continues to correlate with markers of fitness after recruitment in more frequent breeding and higher investment in pups. Though this directional selection is predicted to, if length is heritable, cause long natal body lengths to evolve over time, it remains untested whether the population at large is getting longer as selection pressures from increasing seal density intensify. The 1998-2002 cohorts analyzed here were the first in this population with standard lengths measured in the process of marking for recapture, but lengths were also measured for pups born in more recent cohorts (2014-2016). Preliminary analysis shows that pups born in the 2014-2016 cohorts are 0.66 cm (±0.19 SE, unpublished data) shorter on average than pups born in the 1998-2002 cohorts, so it does not appear that there is any observable response to the selection pressures in pups being produced on Sable Island. This is unsurprising; due to the low survival rate of cohorts recruiting during slowing population growth, the bulk of the reproductive population is still older females born during a period of exponential population growth that experienced very little size selection. So, even if this trait were heritable with selection acting upon it, long reproductive lifespans and low recruitment to the breeding population can drive a slow rate of evolution. If this trait is at stasis, genetic constraints may also prevent the evolution of body size (e.g. a lack of heritable variation, genetic correlations between traits under selection); or, genetic evolution of larger size is occurring but these changes are masked at the phenotypic level by changes in environmental conditions (Pujol et al. 2018, O’Sullivan et al. 2019).

The heritability of a trait may decrease over ontogeny, such that the correlation between offspring traits and parental traits diminish with age or are only apparent in certain stages. Phenotypic variance of body length likely declines with increasing age, due to processes such as directional viability selection that acts in particular over early ontogeny (e.g. survival to reproductive recruitment) and compensatory growth processes. So, females that produce long pups could have been long in early development themselves, but of average length through adulthood. For Sable Island grey seals as well as most other animal populations, very little data exist to determine if there is selection on this trait through later stages of life. Grey seals have particularly high and consistent survival as adults (0.989±0.001 for females aged 4-24, 0.901±0.004 for females aged 25+, den Heyer and Bowen 2017), so directional selection on body length as adults is more likely to act through variation in reproductive performance. Further investigation into changes in size-selective vital rates as the population continues to increase would likely yield important insights into density-related evolutionary changes in long-lived animals.

### Implications for population dynamics

The Sable Island grey seal breeding colony has increased dramatically over the past half century with near maximum population growth of 13% per year between the 1960s and late 1990s (Bowen 2011) and a reduced rate of increase of 4% from 1997 to 2016 (den Heyer et al. 2017, den Heyer et al. 2021). Female grey seals born during a period of exponential population growth in the 1980s and 1990s had apparent juvenile survival probabilities to reproductive recruitment of 0.7–0.8 (den Heyer et al. 2013). By contrast, in the late 1990s to early 2000s, when our study females were born, the population experienced reduced population growth rate as it seemingly approached carrying capacity (Bowen et al. 2007, Bowen 2011, den Heyer et al. 2017, den Heyer et al. 2021), with drastically reduced apparent juvenile survival probabilities ranging from 0.26 to 0.39. Previous analyses suggest a size-selective mortality, with individuals with longer natal lengths more likely to survive to reproductive recruitment (Bowen et al. 2015).

We note that our sampling scheme and modeling framework may yield only a conservative estimate of the relationship between natal length and reproductive performance, as we only included individuals that survived to recruit to the breeding population and (1) observed on the island in at least 2 years and (2) nursed their pup long enough to be recorded by our research teams. These qualifications result in a sample that only explores part of the spectrum of reproductive investment. Inexperienced or low quality mothers may frequently flee or abandon pups, and these reproductive attempts would not be recorded in our observations, though this is not a source of serious bias (Hammill et al. 2017).For these reasons, poor performers are much less likely to be observed, resulting in a larger proportion of high quality females in our sample than present in the Sable Island breeding population.

Our sample of females make up the post-selection distribution of body size, and this study can perhaps be viewed as a lens into the reproductive performance of individuals growing during intense selection pressure and slowing population growth (Coltman et al. 1999, Allen et al. 2008). In addition to our results linking natal size with reproductive success, we found that this sample of females exhibited a slight cost of reproduction not detected when a larger subset of the population was analyzed (Badger et al. 2020), which included females born in the 1960s, 1970s, and 1980s that went through the juvenile stage when population density was much lower. From this, we may infer that ecological conditions during early stages can mediate future trade-offs and shape the natural selection on life history and pace-of-life (Clutton-Brock et al. 1987, Coltman et al. 1999). Intensified competition among these age groups may drive a less favorable energetic trade-off between survival and supporting reproduction for newly recruiting individuals. Alternatively, the discrepancy could be a result of younger, smaller individuals dominating the data set analysed here, versus older, more experienced individuals in the previous analysis. A cost of reproduction may diminish over time, such that it would not be detected in a data set comprised of mainly older individuals. A more formal comparison of reproductive rates among cohort groups incorporating interactive effects of female experience and age is needed to assess how increasing population density may have influenced drivers of life history evolution in grey seals.

The Sable Island grey seal population is still increasing at a rate of 4% per year, and as the Scotian shelf becomes more saturated with seals, juvenile selective survival will likely intensify (Eberhardt and Siniff 1977). If so, the estimated effects of offspring size on maternal fitness may be greater during this period of slowing density dependent population growth, and continue to rise (Clutton-Brock et al. 1987). Long-lived, iteroparous species such as grey seals have high survival rates likely because they are able to vary reproductive effort in response to environmental conditions (Bull and Shine 1979, Allen et al. 2008). Females may be more likely to forego reproduction altogether than support gestation and birth only to provide insufficient or highly variable care to dependent offspring. Mothers may opt instead for fewer, larger offspring more likely to survive and become better reproducers. While this theory is not supported by previous work on lifetime reproductive performance of grey seals using a sample of females born in the 1980s cohorts (Chapter 3), as discussed previously, the nature of life history variation may fundamentally change as competition continues to increase (Clutton-Brock et al. 1987, Coltman et al. 1999, Allen et al. 2008).

### Carryover effects of morphology during early ontogeny

The covariation between natal length and future reproductive performance reported here is more likely to be a carry-over effect from advantages granted to growth and self-maintenance as a juvenile that allow for greater adult performance, rather than a physical trait that is maintained throughout life. Carry-over effects describe how the environment experienced early in life can affect the expression of traits in different life stages and habitats (O’Connor et al. 2014, Moore and Martin 2019). Carry-over effects that occur at the individual level can affect a wide range of fitness parameters, propagating long-term, large scale consequences on population dynamics and composition and so influence multiple levels of biological organization from individuals, populations, and even community structures ((Norris 2005, Betini et al. 2013, O’Connor et al. 2014, Moore and Martin 2019).

Development rarely resets entirely during different environments and life stages–it should be no surprise that these cascading linkages are present across taxa (Moore and Martin 2019) and have been extensively documented over the last two decades. For example, larval experiences of marine invertebrates, such as stress or prolonged swimming time, can have carryover effects on juvenile growth and survival despite the massive tissue reorganization associated with metamorphosis (Marshall et al. 2006). The distribution of body sizes in water pythons reflects stochastic rain-fall induced annual variation in prey availability over the preceding decades (Madsen and Shine 2000), and nonmigratory fish experiencing high stress in spring show differences in growth rates through the summer (O’Connor et al. 2014). Descamps et al. (2008) found that red squirrels (*Tamiasciurus hudsonicus*) that were born under conditions of higher food availability between birth and weaning had a higher reproductive success as adults than those born under lower food availability. A study of *Spartina alterniflora* salt marshes demonstrated that the effects of a single enrichment of nitrogen persisted for up to three years, affecting plant primary productivity long after the nitrogen supplement has been exhausted (Gratton and Denno 2003). Nussey et al. (2007) found that red deer females experiencing high levels of resource competition during early life showed faster rates of senescence as adults. The classic example of a seasonal carry-over effect comes from studies of migratory birds, where birds overwintering in high quality habitats have higher reproductive success in the subsequent breeding season than birds overwintering in less desirable habitats, although the quality of the breeding habitat is the same for both groups (Harrison et al. 2011). Carryover effects provide a substantial source of variation in fitness such that we often cannot understand ecological dynamics of natural populations when only current conditions are observed.

Food availability during the ontogenetic period is a key environmental factor in carry-over effects (Descamps et al. 2008, Harrison et al. 2011), with the ultimate driver being habitat quality, or less commonly reported, intraspecific density. Our results suggest female grey seals experience a carry-over effect of individual characteristics in early ontogeny on future reproductive performance that was likely induced by negative density dependence. Breed et al. (2013) documented that juvenile grey seals may be competitively excluded from key foraging grounds near Sable Island by adult females as the population increases, potentially contributing to the stark decline in juvenile apparent survival to recruitment in the 1998-2002 cohorts (den Heyer et al. 2013). This exclusion may continue into adulthood, such that there is further intense competition to secure ideal foraging grounds, else expend more energy traveling on foraging bouts. It is possible that longer juveniles were more able to compete with adults and secure better foraging habitat, which carried over into reproductive years affecting their reproductive fitness traits (Lloyd et al. 2019). Young grey seals are largely unobserved from weaning until recruitment to the breeding population, so the causal pattern linking natal length to subsequent performance is unknown. Further investigation into the factors influencing juvenile ecology is needed to understand the life-long effects of early ontogeny.

### Conclusions

Using nearly 20 years of reproductive data for a large sample of marked females, we found that size in early stages of ontogeny is positively associated with two measures of reproductive performance later in life. Our findings underscore the multiple lines of evidence before us that have demonstrated that maternal fitness depends on attributes of offspring size and their cascading effects on offspring fitness, and constitute the first documentation of size carry-over effects of early ontogeny on adult performance in pinnipeds, and likely also in marine mammals. In this case, natal size appears to be acting as a carry-over effect coinciding with shifting population dynamics, our findings prompt further investigation into how negative density dependence shapes the evolution of life histories in a long-lived, iteroparous animal. Phenotypic selection across life stages will vary according to how fitness is maximized in a given environment, and will have large-scale consequences in ecological and evolutionary time scales.

